# The effects of a 6-week core exercises on swimming performance of national level swimmers

**DOI:** 10.1101/2019.12.19.882126

**Authors:** Jakub Karpiński, Wojciech Rejdych, Dominika Brzozowska, Artur Gołaś, Wojciech Sadowski, Andrzej Swinarew, Alicja Stachura, Subir Gupta, Arkadiusz Stanula

**Author notes:** These authors contributed equally to this work.

## Abstract

The aim of this study was to assess the impact of a 6-week specialized training program aimed at strengthening core muscles to improve the effectiveness of selected elements of a swimming race in group of elite polish swimmers. Sixteen male national and international level swimmers (21.6 ± 2.2 years) participated in the research. The competitors were randomly assigned to 1 of 2 groups before the data collection process: an experimental (EG, n = 8) and control (CG, n = 8) group. Both groups of swimmers carried out the same training program in the water environment (volume and intensity), while swimmers from EG additionally carried out the specific core muscle training. The task of the swimmers was an individual swim of 50 meters freestyle, during which the kinematic parameters of start, turn and swimming techniques were recorded using the video camera system. In both groups a minor increase in the flight phase was observed during the start (EG=0.06 m, 1.8%; p=0.088; CG = 0.08 m, 2.7%; p=0.013). The time of the distance 5 m after the turn and the recorded average speed of swimming this distance in EG statistically significant improved accordingly 0.1 s (−28.6%; p<0.001) and 3.56 m·s^-1^ (23.2%; p=0.001). In EG, a statistically significant improvement in 50-m freestyle performance was observed by 0.3 s (−1.2%, p = 0.001). The results of the research show that the implementation of isolated strength of the stabilizing muscles seems to be a valuable addition to the standard training of swimmers.

## Introduction

Strength and muscular power are a significant determinants of success in swimming sport. Appropriate training of the abdominal muscles and torso seems to be one of the key elements determining the effectiveness of the training process [1]. The main goal of swimming competition is to overcome the given distance in the shortest possible time, which is achieved, among others, due to proper body positioning in the water and minimizing the resistance [2–5]. Numerous publications show that exercises strengthening the core muscles is an integral part of many training programs [4,6,7]. Increased work of the stabilizing muscles can form the basis for generating more strength through the limbs [8,9]. In many sources the concept of core muscles is expanding, including for rectus abdominal, latissimus dorsi, gluteus maximus or trapezius [6,10]. Proper control of the body position while swimming at a distance, as well as during the start jump and turn, increases efficiency and thus reduces the distance traveled [11].

Appropriate strengthening of the muscles responsible for the correct position of the body is fundamental to the swimming technique [12]. It causes correct positioning of individual body segments, i.e. the head, shoulder girdle, torso, pelvis girdle and legs. Efficiency of those muscle provides a nearly linear arrangement of these segments, thereby minimizing the resistance that puts the body in the water [11,13,14]. The unstable background in which the body of the swimmer is located requires an exemplary core muscle work, and the lack of stable support means that the deficit of one or several muscles can cause huge time losses. In addition to minimizing resistance, the appropriate high and stable body position allows you to optimize the power of your upper and lower limbs [4,15,16].

There is a lot of evidence in the literature on the effectiveness of training on land for the results achieved in swimming [17,18]. In the studies of Patil et al. [4], in accordance with the authors’ expectations, the planned specialist core muscle strengthening training improved the performance of this area (Functional Core Muscle Strength test) and led to significant improvement during a 50 m freestyle.

Also, Gencer [19] observed that during the experiment, there was a significant improvement in time needed to swim the distance of 50 meters by the experimental group in relation to the control group [19]. Similar conclusions came from Gönener et al. [20], who after the tests stated that training with the use of Thera-Band tapes (including engaging core muscles) improves the performance of swimmers [20]. Despite the interest of the authors of numerous publications on the issue of core muscle training and despite the universal recognition that the right strength of stabilizing muscles improves the athletic level of competitors, there are papers that show only the marginal impact of this training on the final sporting success [7,10,19]. The results of many studies also did not show a direct relationship between improving muscular strength on land and improving the results achieved by swimmers in the water [21,22]. Lack of consistent results, their ambiguity make it necessary to continue research on the effectiveness of training on land, including core muscle training and its impact on the sports score.

For this reason, the authors of this study decided to assess the impact of a 6-week specialized training program aimed at strengthening core muscles (SCMT) to improve the effectiveness of selected elements of a swimming race in group of elite polish swimmers.

## Methods

### Participants

Sixteen male national and international class swimmers who are members of the Polish National Swimming Team (seniors and youth) were involved in this research. Participants of the experiment were in the same period of preparation for the competition, i.e. in the sub-period of specific preparation. The competitors were randomized to an experimental group (EG) of 8 swimmers and a control group (CG) also 8 swimmers. Both groups of swimmers carried out the same training program in the water (volume and intensity), while swimmers with EG additionally carried out specialized core muscle training (SCMT), which took place 3 times a week for 6 weeks. The total volume of the applied training program was 288 minutes, of which 216 minutes of effective work with progressing in weekly or biweekly periods. Training performed for the needs of the experiment did not disturb the proper process of preparing swimmers to start in competitions. All participants had current medical examinations and any contraindications to participation in the studies were excluded. None of the swimmers were taking drugs, medication, or dietary supplements known to influence physical performance. During the experiment, the subjects were tested on an equal and balanced diet. Calorific value was selected individually based on the measurement of body mass composition and the volume and intensity of the training program. Swimmers also expressed written consent to participate in the experiment. All competitors were presented with goals, expected duration and course of tests. Athletes were also informed about the possibility of giving up participation in the experiment at every stage. The research project has been approved by the University Bioethics Committee for Research at The Jerzy Kukuczka Academy of Physical Education in Katowice (No. 8/2018). Anthropometric data of competitors are presented in Table 1. The measurement of the body height was made with an anthropometer with an accuracy of 0.5 cm, while the body mass and its composition determined by the method of electrical impedance using the InBody 220 device (Biospace Co. Japan).

**Table 1.** Physical characteristics of participants (mean ± SD).

|  | Experimental group<br>(n = 8) | Control group<br>(n = 8) | *p-value* |
| --- | --- | --- | --- |
| Age (year) | 20.2 ± 1.17 | 20.0 ± 1.9 | 0.606 |
| Weight (kg) | 74.9 ± 10.67 | 75.4 ± 6.27 | 0.926 |
| Height (cm) | 183.0 ± 6.57 | 182.1 ± 3.18 | 0.761 |
| Fat mass (%) | 6.52 ± 3.22 | 10.22 ± 4.92 | 0.154 |
| Lean body mass (kg) | 30.5 ± 5.46 | 29.2 ± 1.50 | 0.577 |
| Fat mass of trunk (%) | 5.75 ± 3.01 | 10.70 ± 5.93 | 0.098 |

### Procedures

The training program lasting six weeks consisted of 18 units of targeted training on land. The duration of the main unit part did not exceed 25 minutes. According to the purpose of the research, the developed training program included exercises involving the core muscles. In the general sense, we interpret them as torso muscles or less recently used the term “body core”. Comparing with each other numerous definitions of this term we find a common denominator, which is the deep muscles that provide stabilization of whole body and the base for functional stability of the lumbar, sacral and iliac area [6,10,12,14]. SCMT consist of four exercises: Flutter kicks (scissors), single leg V-Ups, Prone physio ball trunk extension, Russian twists. Progression consisted of changing the position of the body, adding a motion element, adding an unstable ground and increasing the resistance. The same training units were carried out three times a week. Depending on the exercise, the level of difficulty progressed in weekly or bi-weekly cycles. In the case that the competitor was unable to complete the task with a certain resistance, he returned to the load from the previous microcycle until the end of the duration of the given exercise. All exercises were performed in 4 series in a 40-second work schedule - a 20-second break between sets. A detailed training program is presented in Table 2.

**Table 2.** A brief description of the exercises of SCMT and their progression over 6-week training program.

| Weeks of training | Flutter kicks (scissors) | Single leg V-ups | Prone physio ball trunk extension | Russian twists |
| --- | --- | --- | --- | --- |
| 1 | Arms crossed on the chest | No extra load | Arms crossed on the chest | No extra load |
| 2 | Streamlined position | No extra load | Arms crossed on the chest | No extra load |
| 3 | Arms crossed on the chest + weights on the ankles | Dumbbells in hands | Holding medicine ball | Holding Kettlebell |
| 4 | Streamlined position + weights on the ankles | Dumbbells in hands | Holding medicine ball | Holding Kettlebell |
| 5 | Arms crossed on the chest + weights on the ankles (swimmer performs this progression on wiggle cushion) | Dumbbells in hands + weights on the ankles | Medicine ball trunk extension throw | Holding Kettlebell (swimmer performs this progression sitting on wiggle cushion) |
| 6 | Streamlined position + weights on the ankles (swimmer performs this progression on wiggle cushion) | Dumbbells in hands + weights on the ankles | Medicine ball trunk extension throw | Holding Kettlebell (swimmer performs this progression sitting on wiggle cushion) |

The tests consisted of two stages: preceding the experiment and performed after the experiment. During the research used the same procedure, the same time of day and the order of the athlete. The measurements were carried out on a 25-meter swimming pool (The Jerzy Kukuczka Academy of Physical Education in Katowice) three days before and after the core muscle training was completed. The task of the swimmers was to swim 50 meters freestyle from the starting block under the race conditions. In order to accurately measure the times obtained by the participants, the Omega electronic time measurement system was used (OMEGA S.A., Switzerland). The swimming race was recorded using two digital video cameras (JVC GC-PX100BE, Japan) with a rapid shutter speed (1/1000 s), operating at a sampling rate of 50 Hz. One of the cameras was set 1.5 m above the water at a distance of 2 m from the starting wall perpendicular to the direction of the road traveled by the swimmer in order to register the dive start and enter the swimmer into the water. The second camera was placed 1.5 m above the water exactly in the middle of the length of the swimming pool (12.5 m from the starting wall) to capture the distance swimming. Both cameras were mounted on tripods, positioned on the poolside 0.5 m from the edge of the pool perpendicular to lane 2. In order to register the glide after the turn, the third camera (Sony FDR-X3000, Japan) was placed under water at a distance of 2 m from the turning wall at a depth of 1.0 m at the side wall of the pool basin. These cameras were calibrated using a series of poles of known lengths positioned at specifically known positions throughout the length of the area that the swimmers travelled during each trial. The following parameters of the dive start were analyzed: Entry distance (cm), Time in the air with Take-off (s), Reaction time (s), Time in the air (s), Entry velocity (m·s^-1^), Dive angle (°). The time was measured on a distance of 5 meters after the turn, and then the speed of swimmer after the turn on the distance of the first 5 meters was calculated. Additionally, based on the swimming velocity data and the time of three complete stroke cycles, the stroke rate (SR) (cycles·s^-1^) and the stroke length (SL) (m) were determined (detailed description of the all measured parameters is provided in Table 3). All video files were analyzed by 2 different researchers with the Kinovea (version 0.8.26.) software, which allowed time-motion analysis of registered elements.

**Table 3.** Detailed description of the parameters measured by using the Kinovea software program while swimming 50-meter front crawl.

|  |  |
| --- | --- |
| Entry distance (cm) | Distance from the starting wall to the head entry point. It is considered as the length of the flight and is measured parallel (horizontal) to the water surface. |
| Entry velocity ($\text{m}\cdot\text{s}^{-1}$) | The horizontal velocity of the swimmer travelling through the air during the flight phase before entry into the water (based on the length of the flight phase and the time in the air). |
| Time in the air with Take-Off (s) | The time needed by the swimmer to leave the block following the starting signal. It is considered as the reaction time. |
| Time in the air (s) (Flight phase) | The time from when the swimmer leaves the block to when the swimmer's head enters the water. It is also known as the flight phase. |
| Dive angle (degrees) | The angle at which the swimmer enters the water. It is the angle between the water surface and the central axis of the body at the time when the head touches water surface. |
| Reaction time (s) | The time needed by the swimmer to leave the block following the starting signal. It is considered as the reaction time. |
| Time 5 m after the flip turn (s) | The time needed by the swimmer to reach the 5 m line after the turn. It was measured between pushing off the wall and crossing the 5 m line by the swimmer's head. |
| Average velocity after the flip turn ($\text{m}\cdot\text{s}^{-1}$) | The horizontal velocity, which the swimmer reaches 5 m after pushing off the wall. |
| Swimming velocity ($\text{m}\cdot\text{s}^{-1}$) | The horizontal velocity, which the swimmer obtains during swimming 5 m on the distance. It was measured between 12.5 and 17.5 m during the first and second 25 m. |
| Time of 3 cycles (s) | The time needed by the swimmer to perform 3 strokes. It was measured during the first and second 25 m. |
| Stroke rate ($\text{cycles}\cdot\text{s}^{-1}$) | The time required to perform 3 stroke cycles was measured (in the middle section of the first and the second lap) and then used to calculate Stroke Rate; $\text{SR} = 60 \times 3/\text{tSR}$ (SR- Stroke Rate, tSR- time taken of 3 cycles). |
| Stroke length (m) | The distance covered in one stroke. It is calculated by dividing the swimmer's distance by the stroke rate. The SL calculation is based on the data gathered in 9-m sectors of the distance of 50 m, during both laps (for the first lap between the 15 and 24 meter, and for the second lap between the 40 and 49 meter). |
| Total time for 50 m (s) | The total time to cover the distance of 50 m from the starting signal until the wall is touched by hand of swimmer at the end. |

### Statistical analysis

Means and standard deviations were used to represent the average and the typical spread of values of the all performance variables of swimmers. The Normal Gaussian distribution of the data was verified by the Shapiro-Wilk’s test. Levene’s test for equality of means showed no significant differences in the group variances. A two-way analysis of variance with repeated measures and Bonferroni post hoc test were used to investigate the main effects and the interaction between group-factor (experimental vs. control), timefactor (pre-training vs. post-training), as well as the existence of differences between groups in the initial and final data of all variables.

The magnitudes of differences between results of pre-test and post-test were expressed as relative difference in percent and as a standardised mean differences (Cohen effect sizes). The criteria to interpret the magnitude of the effect sizes were: <0.2 trivial, 0.2—0.6 small, 0.6—1.2 moderate, 1.2—2.0 large and >2.0 very large. Additionally, the absolute and percentage change from pre- to post-test was calculated for all variables for each group.

Statistical power equations to determine a minimum study population at the *p < 0*.*05* level with a power of 0.8 revealed a sample of minimum of 6 subjects in each group. Statistical significance was set at *p < 0*.*05*. To assess the reliability of the digitizing process (intra-observer and interobserver), 6 trials were quantified using intraclass correlation coefficients (ICC), 3 of them were digitized by the researcher and the other 3 by an investigator with experience in the Kinovea software digitization management. All statistical analyses were conducted using Statistica 13.3 (TIBCO Software Inc.).

## Results

Table 4 shows the results of all measurements made before (pre-test) and after training (post-test). In both the EG and CG, after the end of the 6-week training, we can observe an increase in the entry distance during the take-off. In the EG, the improvement was 0.06 m (1.8%, *p = 0*.*088*, ES = *Moderate*), while in the CG the improvement was 0.08 m (2.7%, *p = 0*.*013*, ES = *Moderate*). With the elongation of the flight phase in the EG, a statistically significant increase in of the entry velocity was noted 0.57 m·s^-1^ (4.3%; *p = 0*.*021*; ES = *Small*), accompanied by a statistically significant reduction in the time in the air with take-off by 0.09 s (−9.7 %; *p <0*.*001*; ES = *Very large*). ANOVA revealed a significant interaction (training group × test time point) for the Time in the air with Take-off (F(1.14) = 10.242, *p = 0*.*006*). In both the EG and CG, a statistically significant reduction in reaction time on the starting platform was recorded by 0.09 s (−11.9%, *p = 0*.*001*, ES = *Very large*) and 0.05 s (−6.1%; *p = 0*.*025*; ES = *Moderate*) respectively.

**Table 4.** Pre- and post-training values of performance variables in swimmers. In each data block, the upper row is for the EG, the lower row for the CG.

| Performance variables | Pre-training<br>Mean ± SD | Post-training<br>Mean ± SD | Change<br>$\Delta$ (%) [±95% CI] | *p* | ES / rating | ANOVA (F, *p*) | | | | | |
| --- | --- | --- | --- | --- | --- | --- | --- | --- | --- | --- | --- |
| | | | | | | Time effect | | Group effect | | Time × Group | |
|  |  |  |  |  |  | F | *p* | F | *p* | F | *p* |
| Entry distance (m) | 3.11 ± 0.09 | 3.16 ± 0.08 | 0.06 (1.8%) [-0.01; 0.13] | .088 | 0.66 / Moderate | 13.390 | **.003** | 6.750 | **.021** | 0.390 | .545 |
| | 2.96 ± 0.13 | 3.04 ± 0.12 | 0.08 (2.7%) [0.02; 0.14] | **.013** | 0.65 / Moderate | | | | | | |
| Entry velocity (m·s<sup>-1</sup>) | 12.77 ± 1.65 | 13.34 ± 1.47 | 0.57 (4.3%) [0.11; 1.02] | **.021** | 0.36 / Small | 0.032 | .860 | 0.408 | .533 | 3.006 | .105 |
| | 13.99 ± 2.87 | 13.53 ± 2.81 | -0.46 (-3.4%) [-1.79; 0.87] | .438 | 0.16 / Trivial | | | | | | |
| Time in the air with<br>Take-off (s) | 1.05 ± 0.03 | 0.95 ± 0.05 | -0.09 (-9.7%) [-0.13; -0.06] | **<.001** | 2.14 / V. large | 34.909 | **<.001** | 1.484 | .243 | 10.242 | **.006** |
| | 1.05 ± 0.1 | 1.03 ± 0.08 | -0.03 (-2.7%) [-0.06; 0.01] | .092 | 0.32 / Small | | | | | | |
| Time in air (s) | 0.25 ± 0.04 | 0.24 ± 0.03 | -0.01 (-3.1%) [-0.02; 0.01] | .285 | 0.22 / Small | 0.039 | .846 | 0.762 | .397 | 1.916 | .188 |
| | 0.22 ± 0.05 | 0.23 ± 0.06 | 0.01 (4.4%) [-0.02; 0.04] | .388 | 0.19 / Trivial | | | | | | |
| Dive angle (°) | 40.13 ± 4.36 | 39.75 ± 4.23 | -0.38 (-0.9%) [-3.53; 2.78] | .787 | 0.09 / Trivial | 0.008 | .929 | 1.761 | .206 | 0.405 | .535 |
| | 37.38 ± 3.25 | 37.88 ± 2.95 | 0.50 (1.3%) [-0.27; 1.27] | .170 | 0.16 / Trivial | | | | | | |
| Reaction time (s) | 0.8 ± 0.03 | 0.71 ± 0.03 | -0.09 (-11.9%) [-0.12; -0.05] | **.001** | 2.87 / V. large | 33.727 | **<.001** | 11.525 | **.004** | 2.702 | .123 |
| | 0.83 ± 0.05 | 0.79 ± 0.04 | -0.05 (-6.1%) [-0.09; -0.01] | **.025** | 1.02 / Moderate | | | | | | |
| Times 5 m after turn<br>(s) | 0.43 ± 0.06 | 0.34 ± 0.06 | -0.1 (-28.6%) [-0.12; -0.07] | **<.001** | 1.51 / Large | 41.101 | **<.001** | 4.975 | **.043** | 1.858 | .194 |
| | 0.5 ± 0.11 | 0.44 ± 0.08 | -0.06 (-14.2%) [-0.11; -0.01] | **.026** | 0.65 / Moderate | | | | | | |
| Average velocity 5<br>m after turn (m·s<sup>-1</sup>) | 11.77 ± 1.68 | 15.34 ± 2.8 | 3.56 (23.2%) [2.16; 4.97] | **.001** | 1.54 / Large | 39.577 | **<.001** | 6.129 | **.027** | 9.547 | **.008** |
| | 10.37 ± 2.14 | 11.58 ± 2.11 | 1.22 (10.5%) [0.1; 2.33] | **.037** | 0.57 / Small | | | | | | |
| Stroke rate<br>(cycles·s<sup>-1</sup>) | 1.02 ± 0.08 | 1.03 ± 0.08 | 0.02 (1.5%) [-0.01; 0.04] | .242 | 0.19 / Trivial | 1.802 | .201 | 3.357 | .088 | 0.628 | .441 |
| | 0.97 ± 0.04 | 0.97 ± 0.05 | 0 (0.4%) [-0.01; 0.02] | .633 | 0.09 / Trivial | | | | | | |
| Stroke lenght (m) | 1.63 ± 0.15 | 1.58 ± 0.16 | -0.05 (-3.5%) [-0.12; 0.01] | .091 | 0.36 / Small | 3.496 | .083 | 0.063 | .805 | 3.238 | .094 |
| | 1.59 ± 0.06 | 1.59 ± 0.08 | 0 (-0.1%) [-0.03; 0.02] | .924 | 0.01 / Trivial | | | | | | |
| Total time 50 m (s) | 25.24 ± 0.35 | 24.94 ± 0.49 | -0.3 (-1.2%) [-0.43; -0.16] | **.001** | 0.71 / Moderate | 8.890 | **.010** | 15.13 | **.002** | 0.580 | .458 |
| | 26.82 ± 1.09 | 26.64 ± 1.19 | -0.18 (-0.7%) [-0.53; 0.18] | .274 | 0.16 / Trivial | | | | | | |

Note: CI – confidence interval; ES – effect size: <0.2, trivial; 0.2—0.6, small; 0.6—1.2, moderate; 1.2—2.0, large; >2.0, very large; $\Delta$ (%) – Absolute and percentage of change from pre- to post-test; *p* – *p*-Value

At the end of the experiment, the time of the distance of 5 m after the turn and the recorded average speed of swimming this distance in both the EG and CG have improved. In the EG, the above-mentioned elements of the swimming race were significantly improved by 0.1 s (−28.6%, *p <0*.*001*; ES = *Large*) and 3.56 m·s^-1^ (23.2%; *p = 0*.*001*; ES = *Large*), respectively, while in the CG, these parameters improved by 0.06 m·s^-1^ (−14.2%, *p = 0*.*026*, ES = *Moderate*) and 1.22 m□s^-1^ (10.5%, *p = 0*.*037*, ES = *Small*) respectively. ANOVA revealed of a significant interaction for the Average velocity 5 m after turn (F(1.14) = 9.547, *p = 0*.*008*).

The result of all observed changes was the value of the last of the tested parameters - the time of covering the distance of 50 m with a front crawl. In EG, a statistically significant improvement in athletic performance was observed by 0.3 s (–1,2%; *p=0,001*; ES=*Moderate*), while swimmers with CG had a statistically insignificant improvement in athletic performance by 0,18 s (–0.7%; *p=0*.*274*; ES=*Trivial*).

## Discussion

In this article, it was hypothesized that in a selected group of swimmers in the age category of a senior and an adolescent, strengthening the core muscles will positively affect the effectiveness of the studied elements of the swimming race at a distance of 50 m, which may lead to an improvement in the sports result.

Both in the EG and CG, the parameter of the entry distance improved, which may indicate a positive aspect of the training carried out by the competitors in a given period. In addition, it is worth noting that specialist core muscle training did not disturb the value of this parameter, further reducing the start jump time (parameter improvement by 0.09 s, ES = *Very large*), which is the resultant of the reaction time and flight phase time measured until the swimmer touches the water surface with the head. The value of this parameter among EG athletes was statistically significantly lower (p <0.001), while the improvement that followed was higher than in CG. It is worth noting that in the EG there was a statistically significant increase in the speed of entry of the swimmer to water (4.3%, ES = *Small*), in contrast to CG, in which the regression of the analyzed velocity (−3.4%) occurred.

As the research results show, swimming start directly affects the sports level, depending on the competition, and especially the distance covered, it is 0.8% of the time obtained at 1500 m and 26.1% at 50 m (freestyle) [23,24]. According to the assumptions of this experiment, one of the analyzed parameters was the start jump, which can be divided into three stages: on the start block, flight and underwater phase [24]. In this work, the first two were analyzed and it is worth noting that under the influence of SCMT, the swimmers studied improved the reaction time. This studies did not include the assessment of the reaction time to the starting signal as a skill determined by individual neuromuscular (non-mechanical) skills, and the tested time (on the starting platform) was measured from the moment of the start signal to the moment of loss of contact between the foot and the starting block [25,26]. It is worth noting that under the influence of SCMT, the swimmers significantly statistically improved the reaction time (*p = 0,001*; ES = *Very large*).

These results are consistent with the work of Rejman et al. [25], in which the time on the starting block was shortened due to the six-week plyometric training, and the speed of the swimmer achieved during the flight phase increased (0.71 m·s^-1^), which may be related with improvement in the lower limbs power [25]. Although there is no statistically significant increase in swimmer entry velocity to water, it is worth noting that in the EG there was an improvement in this parameter as opposed to the CG, in which there was a statistically insignificant regression of the analyzed velocity. It seems that under the influence of core muscle training the integration of the muscles of the lower and upper limbs and torso improved, which translated into a more efficient transfer of energy from the lower limbs to the body, and further to the arms and thus more efficient (faster) torpedo (starting) position [8,15].

Many studies show the importance of the flight phase, the maximization of which, combined with the appropriate entry to the water, allows the swimmer to achieve higher speeds during the underwater phase [27,28]. The distance of the flight phase is a very important parameter of the effectiveness of the swimming race, because the body travels much faster in the air than in the water [29]. In Ray and Young [30] studies, ground resistance training did not affect the distance of the flight phase during the starting jump, which may be related to its specificity. In the studies conducted by authors in both EG and CG, there was a statistically significant improvement in the length of the flight phase parameter, which may indicate a positive aspect of the training carried out by the swimmers in the given start-up preparation period. It is worth noting that specialist core muscle training did not disturb the value of this parameter, and additionally shortened the start-up time, which is the resultant of the reaction time and flight phase time measured until the swimmer touches the water surface with the head. The glide speed after the start jump is highly dependent on the time of entry into the water, swimmer position, direction and depth of entry [31,32]. In studies based on a correlation analysis, it was determined that there is a strong relationship between “take-off horizontal velocity and time on Block” and the time obtained by competitors at the initial distance of 15 meters [27,33]. Increasing the take-off horizontal velocity should cause the swimmer to enter the water at a smaller angle. In the conducted studies, in the EG group, an improvement of the flight phase velocity and decrease of the swimmer entry angle (statistically insignificant) was observed, while in the CG group both parameters did not improve. According to other studies, it can be presumed that the wrong position of entry into the water, despite the appropriate speed of the starting jump, will not translate into the speed that the swimmer will reach during the underwater phase [26].

In both analyzed groups, a decrease in swimming time in the first 5 m distance after the turn was observed, and the decrease in this value was statistically significant in the EG (time better by 28.6%, *p<0,001*; ES=*Large*). What’s more, it significantly influenced the next analyzed parameter, the speed of the swimmer crossing the distance of 5 m after the return from the turn wall, this value improved by 23.2% (*p=0*.*001*; ES=*Large*). There are very few researches in the literature investigating the effectiveness of swimming turn, especially tumble turn, due to, among others, lack of appropriate technologies. The technical element that is a swimming turn is complicated due to the environment in which it takes place, multilevel and multi-axis movement and the number of involved body segments [34]. It is undeniable that a properly performed turn contributes to the improvement of the total time of swimming the distance [34]. It is known that a slight improvement in the components of the turn can improve the effectiveness of swimming the total distance. One of the elements of the turn is the glide, which may depend on the push-off and the proper position of the swimmer’s body [4,11]. It seems that the decrease in the time of crossing the first meters after the turn may significantly affect the final time of the distance. In the EG, an increase in the stroke rate [SR] by 1.5% was observed; ES = *Trivial*), as well as shortening the swimming stroke length [SL], which in competitors performing core muscle training, decreased by 3.5% (*p = 0*.*091*, ES = *Small*). There were no significant changes among CG competitors. The increase in the speed of swimming may be caused by an increase in stroke length with a simultaneous drop in the stroke rate, but it can also be achieved by extending the swim stroke length without changes in attendance [35]. In the work of Patil et al. [4] there were no statistically significant changes in the stroke rate and stroke length under the influence of core muscle training. The lack of similar results of various studies can be explained by among others another preparatory period in which experiments were carried out. In addition, many authors have determined stroke rate and stroke length as a factor of swimming performance, which is associated mainly with strength and muscular power [36]. It is also worth to be noted that the competition on mentions distance researched by the authors for the needs of this study is characterized by high dynamics. It seems that the desired effect of directed stabilization muscle training will be observed at a longer distance, eg 200 m, where the correct position of the swimmer’s body seems to be crucial, and thus the stroke length may be longer.

The resultant of all observed changes was the value of the last of the tested parameters - the time of crossing the distance of 50 m. In EG, a statistically significant improvement in athletic performance was observed by 1,2% (*p=0,001*; ES=*Moderate*). During the final test, swimmers in CG also achieved a better result, but this improvement was not statistically significant. On the basis of the available literature, a rational explanation of this issue may be the fact of increased core muscle activity, which allows for a more effective transfer of strength between the limbs, and additionally, for maintaining the body in a streamlined position [11,37]. There are many papers on the impact of land training on performance in swimming sport, however, the results of these studies are not consistent. For example, Tanaka et al. [37] suggest that the increase in strength through a resistance land training does not affect the swimmer’s driving force in the water, and therefore, does not improve swimming performance [37–39]. A large progression was observed in the study of Weston et al. [10], however, it may be caused by a much longer period of the specialist training program, as well as a younger research group. Another study found an improvement in central stabilization that did not translate into swimming efficiency [4]. However, there are a numerous studies proving the positive impact of training on land on the results of swimming, and the recorded progression of results oscillate between 1.3% and 4.4% [12,40]. The results obtained by the authors of the above studies are similar to the results of the work of Weston et al. [10] in which, as a result of twelve weeks of training involving core muscles, a 2% improvement in sprinting distance was observed. Patil et al. [4] also noted a statistical progression of the sport result after a six-week training to strengthen the stabilizing muscles.

It is worth noting that in these studies the improvement in the efficiency of individual swimming elements translated into the final sports result, which is the time of crossing the distance of 50 m. The authors noted a progression of 1.07%, which at the sprint distance seems to be an extremely important improvement, which in direct sports competition can guarantee final success. The CG did not improve the time.

## Conclusions

The present study was attended by a group of selected elite swimmers, who completed a specially designed training program aimed at improving strength and endurance of core muscles. The research results suggest that the implementation of isolated training to strengthen the stabilizing muscles seems to be a valuable addition to the standard swimmers training [10]. Based on the conducted experiment, it can be concluded that the described training affects the efficiency of swimming on a short distance. It is especially worth noting that in these studies, the improvement in the efficiency of individual swimming elements has translated into the final sports result, which is the time to swim the distance of 50 m. The authors noted a statistically significant progression of the result, which seems to be fundamental at the sprint distance. In direct sports competition, even a slight improvement in time may guarantee final success. On the other hand, the similar results of this and other experiments indicate the need to continue research in the field of widely understood land training for swimmers, especially the need to strengthen core muscles. Future experiments should also be enriched with EMG tests showing proper and conscious tensioning of stabilizing muscles.

